# Radiologic Evaluation of the Influence of Cleft Type on Nasal Dorsum Growth

**DOI:** 10.1101/806406

**Authors:** Lingling Pu, Renkai Liu, Bing Shi, David W Low, Chenghao Li

## Abstract

**Purpose:** The study was designed to evaluate whether intrinsic morphological characteristics of the nasal dorsum are affected by cleft type, specifically cleft lip only (CL) and cleft lip with cleft palate(CL/P).

**Methods:** 576 cleft patients (278 CL only, 298 CL/P), and 333 individuals without orofacial clefts were retrospectively enrolled. Lateral cephalometric radiographs of all individuals were taken to evaluate the nasal length and nasal dorsum height. Dunn’s test was used to analyze the difference (*p* < 0.001).

**Results:** In CL and control, the angulation of the nasal bone and nasal dorsum increase by age similarly (5y-18y*, p*>0.05). In CL, the total dorsal length is significantly shorter (5y-18y, *p*<0.001). Although the upper nasal dorsum is similar (except in 5y-6y), the lower nasal dorsum is shorter (5y-18y, *p*<0.001).

In CLP, there is no significant difference in the nasal bone angle compared with controls between 5y-7y. However, it develops insufficiently as children grow (8y-18y, *p*<0.001). The nasal dorsum angle is notably smaller (5y-18y*, p*<0.001). Nasal bone length is not significantly different from control at all stages except during ages 11y-13y (*p*<0.05). Total nasal dorsal length is similar to the control at skeletal maturity (17y-18y, *p*>0.05), although it is shorter during 8y to 16y (*p*<0.05). The upper nasal dorsum is overdeveloped (14y-18y, *p*<0.05), whereas the lower nasal dorsum is underdeveloped (5y-18y, *p*<0.001).

**Conclusion:** CL inhibits the growth of nasal dorsum length, leading to short nose deformity. CL/P patients are prone to saddle-nose deformity because of the diminished nasal height (decreased nasal angle).

## Introduction

Cleft lip is frequently accompanied by nasal deformities. The congenital anatomic deficiency or aberrancy, potential changes related to growth, the cleft itself, and even scarring from previous procedures are the main factors which lead to a wide variability in secondary cleft nasal deformities and the complexity of surgical techniques over the past few years^1^. Subsequently, secondary surgery for the cleft nasal deformity undeniably presents a formidable challenge to the plastic surgeon, and the results are not as ideal as expected due to lack the comprehensive inward characters that hidden under complex deformed manifestations.

Due to its central location, the nose plays a prominent role in facial aesthetics ^2^. How does one distinguish the different factors that contribute to the cleft nasal deformity, including cleft type, intrinsic potential changes, or surgical damage? Nowadays, surgeons mainly define the cleft nasal malformation with regard to the alar base, columella, nostril, nasal tip, nasal floor, and nasal septum, attaching more importance to the dysmorphia of the nasal tip ^3, 4^. However, because of the complexity of this anatomic structure, it is so difficult to define the key factors. As an important part of nose, the nasal dorsum plays a major role in nasal and facial harmony ^5^.Analysis of rhinoplasty results has shown that even slight differences in nasal shape can transform the look of an individual’s face ^6^, the key point being that one might be able to distinguish the effect of different cleft types on nasal dorsum deformity, because of its simpler anatomic structure.

Based on the above reasons, this study focused on evaluating morphologic characteristics of the nasal dorsum, to analyze the role of cleft type on nasal dorsum growth in cleft lip patients with and without cleft palate. The study population included patients with cleft lip only (CL), cleft lip and cleft palate (CL/P), and healthy individuals. The soft and hard tissue of the nasal dorsum was analyzed through lateral cephalometric radiographs to obtain objective data of the hard and soft tissue morphology of the three groups in different ages, then compared. Lateral cephalometric radiographs of all individuals were taken to evaluate the nasal length, including the length of the nasal bone, the nasal dorsum, upper nasal dorsum and lower nasal dorsum. The angulation of the nasal bone and the nasal dorsum were evaluated as the indexes of nasal dorsum height. The results indicate that CL inhibits the growth of nasal dorsum length, leading to a short nose deformity, while CL/P tends to result in a saddle nose because of decreased nasal height. These findings help characterize nasal dorsum development, provide comprehensive characteristics of the secondary nasal deformity in cleft patients, and potentially improve the outcome of secondary reconstructive surgery.

## Methods

### Ethics statement

Samples were collected in accordance with the guidelines of The West China Hospital of Stomatology Institutional Board(WCSHIRB). The experimental protocol was approved by local ethics committee (WCSHIRB, Sichuan University, China). Informed consent was obtained from all subjects or, if subjects are under 18, from a parent and/or legal guardian.

### Sample

The study sample comprised a total of 909 Chinese children aged between 5 and 18 years at the West China Hospital of Stomatology, Sichuan University, Chengdu, China, between 2011 and 2016, who were divided into CL only, CL/P, and a control group. The CL group was composed of 278 children with cleft lip, and the CL/P group was comprised of 298 children with combined cleft lip and palate. They were non-syndromic and had no other congenital anomalies. Following our cleft center protocol, the CL group was treated with a modified Millard technique at 3-6 months ^7^. The CL/P group underwent the same lip repair technique, and then underwent a Sommerlad palatoplasty at 9-12 months. None received any other secondary surgery such as lip revision, fistula repair, rhinoplasty, or orthopedic treatment except for bone grafting at 9-12 years of age. The control group was composed of 333 healthy children without cleft or any other congenital anomalies of the same age range as the CL and CL/P groups randomly chosen from Department of Orthodontics in West China Stomatology. These children underwent simple orthodontic treatment and had normal skeletal relationships, symmetric faces, and no history of craniofacial surgery. All groups were divided by age from 5 to 7 years, 8 to 10 years, 11 to 13 years, 14 to 16 years, and 17 to 18 years (Table 1).

**Table 1.** Sample distribution by age.

|  | 5-7y |  |  |  | 8y-10y |  |  |  | 11y-13y |  |  |  | 14y-16y |  |  |  | 17y-18y |  |  |  |
| --- | --- | --- | --- | --- | --- | --- | --- | --- | --- | --- | --- | --- | --- | --- | --- | --- | --- | --- | --- | --- |
|  | N | mean | SD | p | N | mean | SD | p | N | mean | SD | p | N | mean | SD | p | N | mean | SD | p |
| CL | 44 | 5.9 | 0.80 |  | 35 | 8.9 | 0.73 |  | 67 | 12.0 | 0.80 |  | 78 | 15.3 | 0.75 |  | 54 | 17.5 | 0.50 |  |
| CLP | 68 | 6.1 | 0.84 |  | 63 | 8.9 | 0.84 |  | 63 | 11.9 | 0.74 |  | 58 | 15.1 | 0.83 |  | 46 | 17.5 | 0.50 |  |
|  |  |  |  | 0.17 |  |  |  | 0.79 |  |  |  | 0.90 |  |  |  | 0.17 |  |  |  | 0.91 |
| Control | 64 | 6.1 | 0.77 |  | 74 | 9.0 | 0.83 |  | 75 | 12.0 | 0.82 |  | 73 | 15.0 | 0.82 |  | 47 | 17.5 | 0.51 |  |
| Total | 176 | 6.1 | 0.81 |  | 172 | 9.0 | 0.81 |  | 205 | 12.0 | 0.79 |  | 209 | 15.1 | 0.80 |  | 147 | 17.5 | 0.50 |  |

Table1 shows the sample distribution by age. There is no difference in sample distribution by age in each Group

### Cephalometric analysis

Because our study was a retrospective case-control study using the archive, all the patients took lateral cephalometric radiographs just for clinical needs. Lateral cephalometric radiographs were taken for each subject under standardized conditions with the head oriented along the Frankfort horizontal plane (FH) parallel to the floor.Subjects were asked to relax their lips in a resting position, and to place their teeth in centric occlusion. An EASYMTIC 3298-125 Cephalometry X-ray machine (Chemetron Co., Chicago, IL, USA) was used for all subjects. In order to reduce the influence of maxillary hypoplasia, a reliable craniofacial reference plane “Sella–nasion S-N” was selected, and maxillary and nasal parts were separated by a vertical line through point nasion. Three hard and three soft tissue landmarks were digitized by one observer. Anthropometric landmarks on the nose were defined ^8, 9^. Nasal Dorsum was measured by its length and angular of the hard and soft tissue. Fig.1 shows the landmarks that were used in the cephalometric analysis directly and indirectly, including four linear measurements and two angular measurements. The angulation of the nasal bone and the nasal dorsum were evaluated as the indexes of nasal dorsum height. The parameter measurements are shown in Fig. 2. Each parameter was measured three times repeatedly and the mean was recorded, P25 (First Quartile), and P75 (Third Quartile).

**Fig 1.**
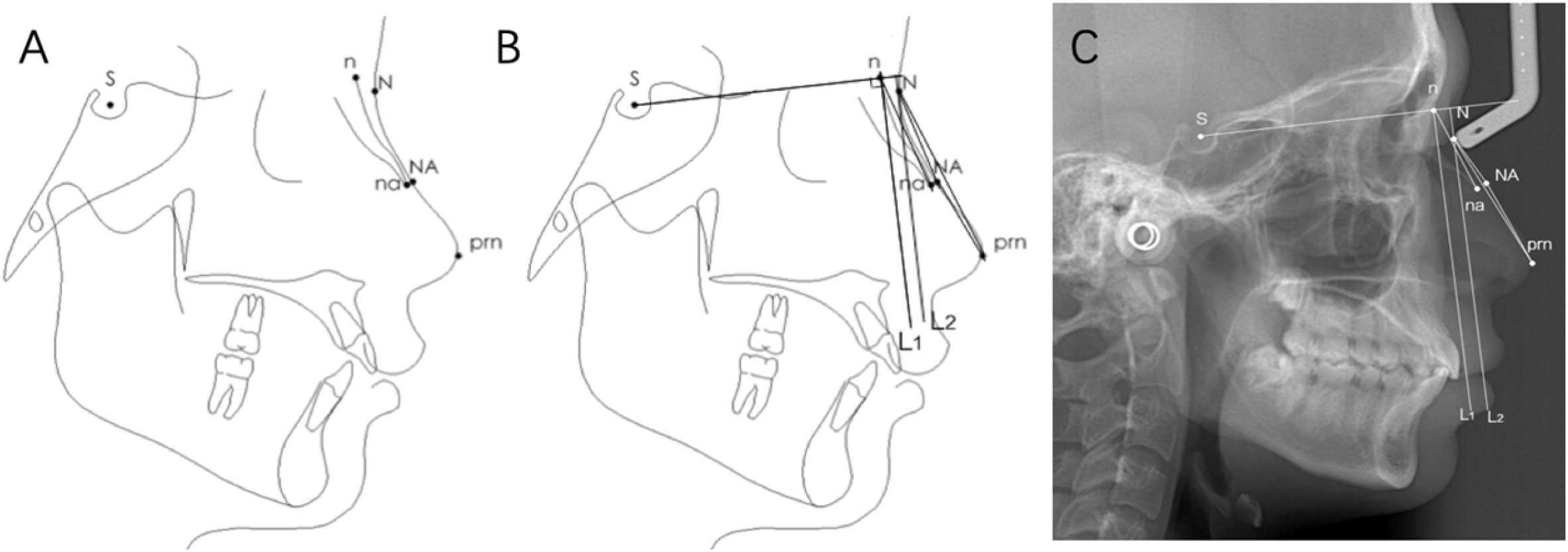
The profile cephalometric radiographs and the cephalometric radiographs. (A)Reference points on the profile cephalometric radiographs. S =sella, the center of sella turcica; n =nasion, junction of frontal, maxillary, and nasal bones; N=soft nasion, closest point on soft tissue outline from hard tissue nasion; na=nasale, point at the most anterior inferior part of the nasal bone; NA= soft nasale, closest point on soft tissue outline from hard tissue nasale; Prn=pronasale, most anterior point on the contour of nose. (B) Measurements of angles and lines on the profile cephalometric radiographs. L1=vertical line of S to n, n is the foot point; L2=parallel to L1; na-n-L1(degrees):angulation of nasal bone; the angle between na-n-L1; prn-N-L2(degrees): angulation of nasal dorsum; the angle between prn-N-L2; n-na(mm):length of nasal bone, from the nasion to nasale; N-prn(mm):length of nasal dorsum, from soft-tissue nasion to pronasale; N-NA(mm):upper nasal dorsum; NA-prn(mm):lower nasal dorsum. (C) Measurements of angles and lines on the lateral cephalometric radiograph.

**Fig. 2.**
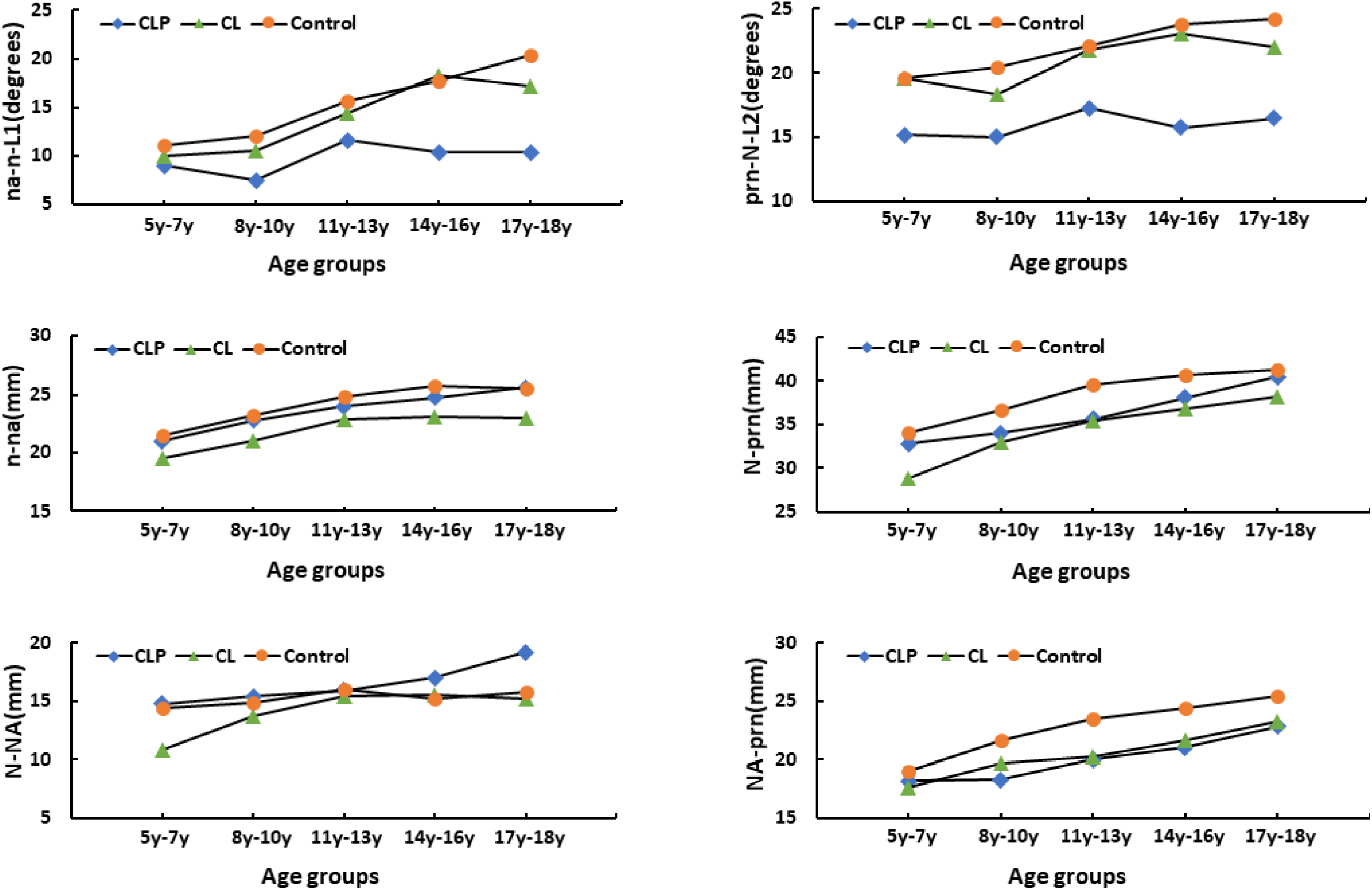
(A) Angulation of the nasal bone changes by years in different groups;(B) Angulation of the nasal dorsum changes by years in different groups;(C) Length of the nasal bone changes by years in different groups;(D) Length of the nasal dorsum changes by years in different groups;(E) Length of the upper nasal dorsum changes by years different groups;(F) Length of the lower nasal dorsum changes by years in different groups.

### Statistical analysis

All statistical analyses were performed with Statistical Package for Social Sciences (SPSS) software version 22.0. ANOVA analysis was used to determine the differences of age distribution in the three groups. Differences in the cephalometric results among the three groups were based on Dunn’s test. The significant difference was defined at 95% level.

### Reliability

To calculate the method error, 100 cephalograms were selected randomly and measured twice, to examine the intra-class correlation coefficient (ICC) ^10^, The ICC results for test-retest reliability ranged between 0.90 and 0.98, suggesting dependable reliability and reproducibility of the adopted measuring strategy. (Table 2)

**Table 2.**
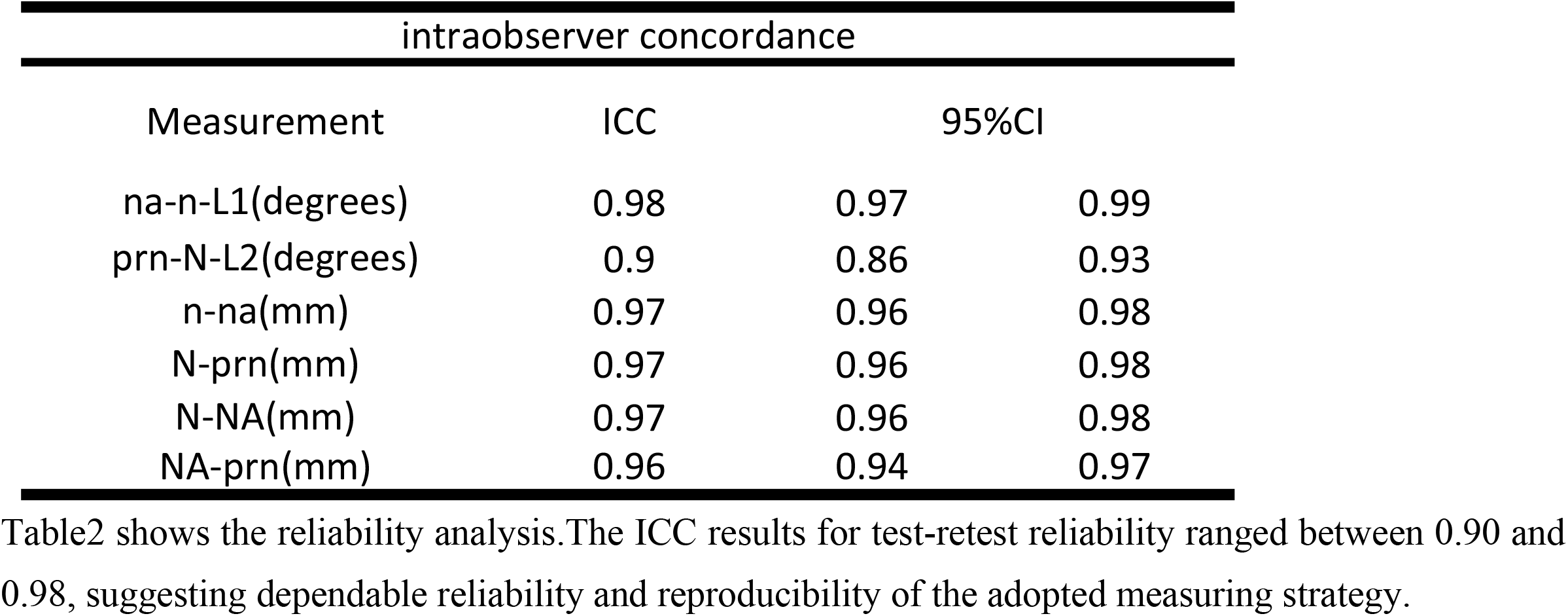
Reliability analysis.

## Results

There was no significant difference in the age composition of CL, CL/P and Control groups. Nasal morphology in three groups was comparable (Table 3). Fig 2 shows the growth tendency of each index in the CL, CL/P and Control group. Fig 3 is the nasal profile map of three groups in 17y-18y.

**Fig. 3.**
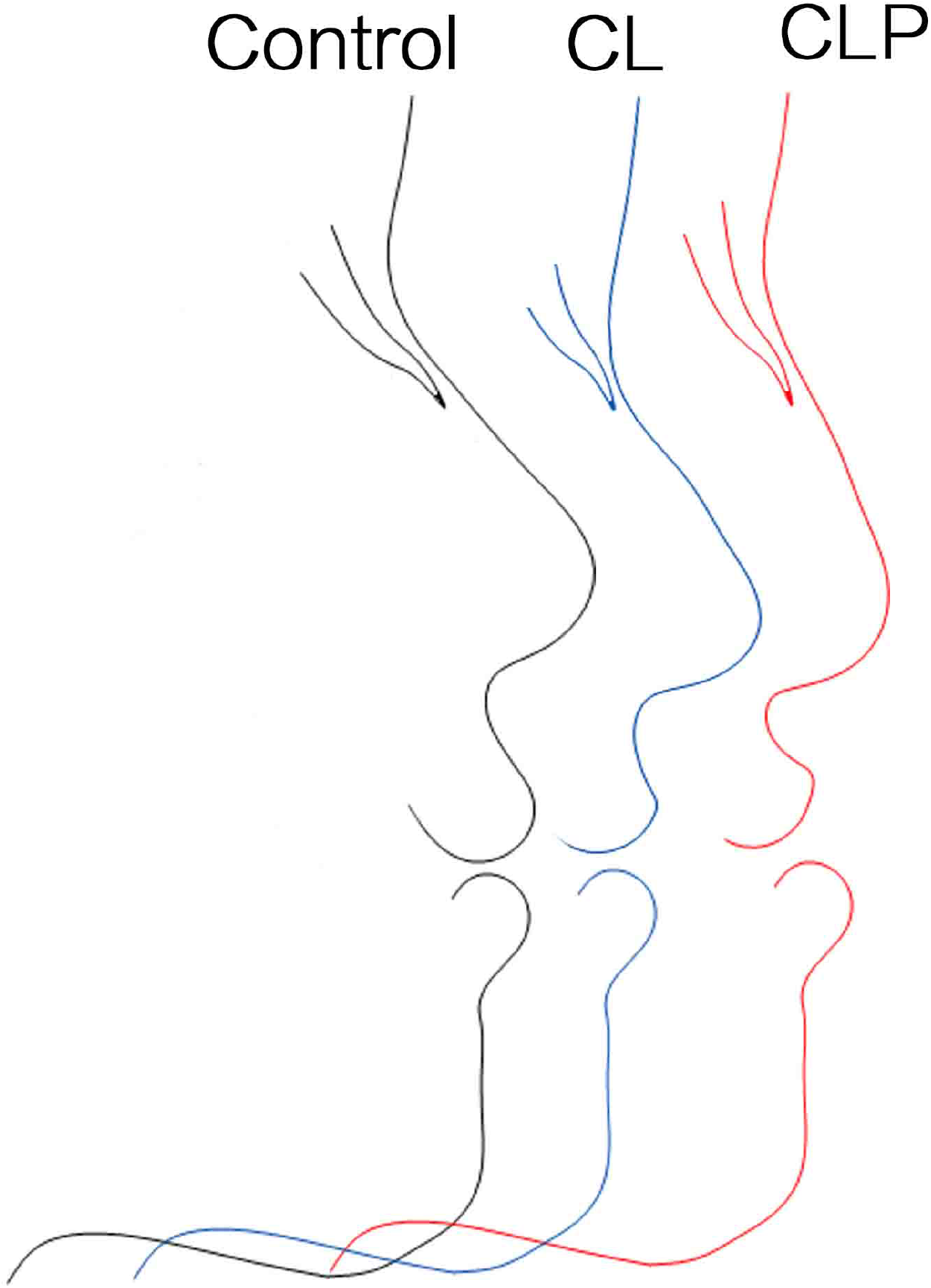
The characteristic features of 17y-18y group determined from our results. The CL patients have a shorter nasal dorsum. The CL/P patients have a flatter nasal dorsum.

**Table 3.**
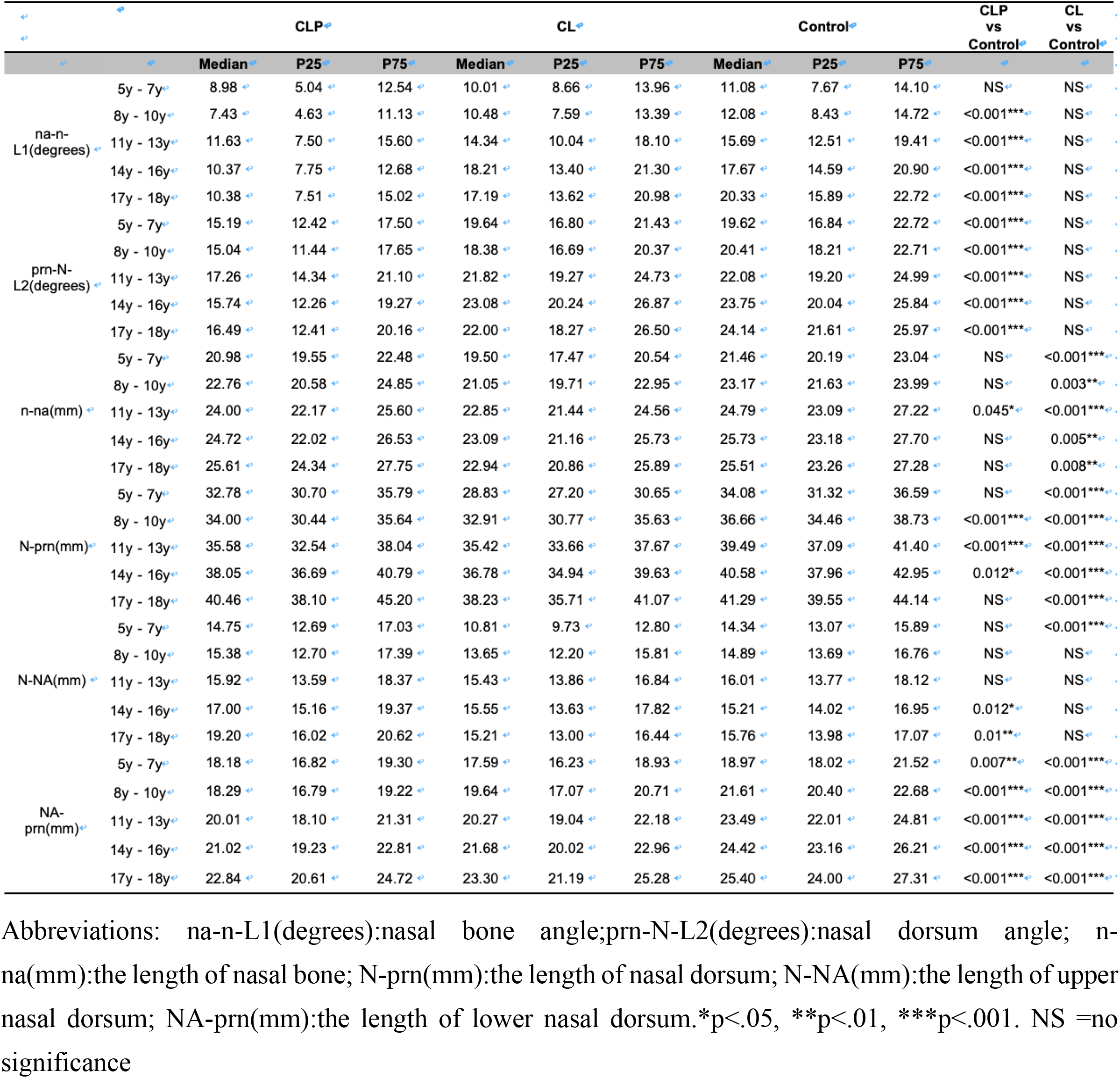
Statistical descriptions of the angle of nasal bone, the angle of nasal dorsum, the length of nasal bone, the length of nasal dorsum, the length of upper nasal dorsum and the length of lower nasal dorsum by ages and results of Dunn’s-test between CLP, CL and Control separately

### 1. CL patients show shorter nose and normal nasal angulation

In CL, compared with Control, the angulation of the nasal bone and nasal dorsum increase similarly by age (5y-18y*, p*>0.05), (Fig 2a, 2b), while the total dorsum length is significantly shorter (5y-18y, *p*<0.001), (Fig 2c, 2d). In CL, the upper nasal dorsum is similar to Control (except in 5y-6y), (Fig 2e), but the lower nasal dorsum is shorter (5y-18y, *p*<0.001), (Fig 2f).

### 2. CL/P patients have flatter angulation, but normal-length nose

In CL/P, there is no significant difference in the angulation of the nasal bone compared with Control in the 5y-7y age range. However, it develops insufficiently as children grow (8y-18y, *p*<0.001),(Fig 2a). The angulation of the nasal dorsum is notably smaller than that in non-cleft children (5y-18y*, p*<0.001),(Fig 2b). Nasal bone length is not significantly different from the control peers at all stages except the peers between 11y and 13y (*p*<0.05),(Fig 2c). At skeletal maturity, the nasal dorsum grows as long as the Control group (17y-18y, *p*>0.05), although it is shorter prior to that (8y-16y, *p*<0.05),(Fig 2c).

The upper nasal dorsum is overdeveloped (14y-18y, *p*<0.05) while the lower nasal dorsum is underdeveloped (5y-18y, *p*<0.001),(Fig 2d, 2e), the net effect being a total dorsum length similar to controls at skeletal maturity.

## Discussion

Our previous works have analyzed the craniofacial and soft tissue morphology of patients with CL/P and CP ^11, 12^. As mentioned earlier, this study was designed to distinguish cleft type factors associated with nasal dorsum deformity. With secondary rhinoplasty mainly aimed at patients with CL regardless of CP, and to eliminate the influence of maxillary retrusion after primary repair on nasal dorsum shape, our study encompassed patients with CL, CL/P, and healthy peers.

In this study, in CL only patients, we found the nasal bone and nasal dorsum were significantly shorter (Fig. 2c, 2d). This is consistent with previous findings: prior studies have demonstrated that in fetuses, newborns, children, and male adults, compared with normal peers, patients with isolated cleft lip had a significantly shorter nasal bone ^13, 14^. Nasal tip position is one of the important indicators for the measurement of the length of nasal dorsum. Compared with healthy ones, CL (with or without CP) showed significant upward deviation in the nasal tip, suggesting that CL patients have a congenital tendency toward a short nose ^15^, but previous researchers have not discussed the nasal features of CL and CL/P separately. Meanwhile, few studies have presented the data from our research, specifically that the nasal dorsum height in CL does not differ from controls (Fig 2a, 2b).

The nasal bone length in CL/P was not significantly different compared with control (Fig. 2c), which was consistent with the conclusion of other authors who have demonstrated that patients with CP +/− CL showed normal length of the nasal bone^13, 14, 16^. The length of the nasal dorsum of CL/P showed growth retardation, which was significantly shorter than that of controls in 8y-16y, but there was no significant difference in the 5y-6y and 17y-18y groups (Fig. 2d). Moreira, et al. ^17^ analyzed the lateral cephalometries of 70 white children with CL/P who had undergone primary operation and found that they had similar nasal dorsum length. But Ferrario, Chiarella, Claudia, Laura, Armando ^18^ found that CL/P patients have a shorter nasal dorsum. Considering the limit of the above sample sizes, we tried to resolve this issue by enrolling a larger sample size, and analyzed the development of hard and soft tissues of the nose in different groups in detail. Additionally, in order to reduce the influence of maxillary hypoplasia, we presented a new evaluation method by utilizing Bookstein, FL and Nadia, H’s design. The results showed nasal dorsum height in CL/P to be lower than controls (Fig 2a, 2b). Therefore, in addition to maxillary hypoplasia, CL/P also demonstrates a flatter nasal dorsum.

CL and CL/P both had a shorter lower nasal dorsum than control (Fig. 2f). Shape changes of the nasal dorsum are most closely related to angulation changes of the lower dorsum ^19^, which may emphasize the malformation of the lower nasal dorsum leading to the whole nasal deformity. The upper nasal dorsum of CL/P was longer than controls, and the difference in 14y-18y was statistically significant (Fig. 2e). However, there was no significant difference in the length of the entire nasal dorsum in 17y-18y (Fig. 2d). We conclude that overdevelopment of upper nasal dorsal length in CL/P compensates for hypodevelopment of lower nasal dorsal length, and the net result leads to a similar length of the entire nasal dorsum compared with controls when growth is completed. However, the developmental mechanism of the upper nasal dorsum deserved further elucidation.

The deformity of the nasal dorsum in CL/P is mainly due to underdevelopment of the height of the hard and soft tissues of the nose, and patients with CL present a shorter nose instead of a flatter one. The characteristic features of the nose for CL, CL/P, and control groups in 17y-18y are shown in Fig.3, providing a basis for a specific approach to secondary rhinoplasty. Most of all, our study confirms that different types of clefts indeed influence the features of nasal dorsum deformity. A flatter nasal dorsum contributes to a flatter profile in patients with CL/P. Hence according to our results, for CL, secondary rhinoplasty should lengthen the nasal dorsum, and for CL/P, the aim of surgery should make the nose more prominent.

## Conclusion

In this study, we evaluated whether morphologic characteristics of the nasal dorsum were affected by cleft types in different ages after primary operation. The results indicate that isolated CL inhibits the growth of nasal dorsum length which leads to a short nose deformity, while the C/LP patients tends to develop a saddle nose because of reduced dorsal angulation which leads to a decreased nasal height.

## Contributions

All authors contributed extensively to this work. Conceived and designed the experiments: L.L.P. and C.H.L. Performed the experiments: L.L.P. and R.K.L. Analyzed the data: L.L.P. and B.S. Interpreted the data: L.L.P. and C.H.L. Wrote the paper: L.L.P., D.W.L, and C.H.L. All authors reviewed the manuscript.

